# A colony-level optimization model provides a potential mechanism for the evolution of novel castes in eusocial ant colonies

**DOI:** 10.1101/2020.10.08.331207

**Authors:** Suryadeepto Nag, Ananda Shikhara Bhat

## Abstract

Ant species often have multiple morphologically distinct ‘castes’ within a single colony. Given that most of these castes are involved in non-reproductive tasks, and since such individuals thus never reproduce, the question of how ant castes can evolve is a non-trivial one. Over the years, several models have been proposed in order to explain the evolution of castes in ant colonies. Here, we attempt to answer this question using an economics-based approach, developing an optimization model that implements adaptation and selection at the colony level. We argue that due to the nature of ant colonies, selection is shifted to the group level, and, due to this, individual ants are sheltered from negative selection. We show that our framework can explain the evolution of novel castes in ant colonies, and discuss the novelty of our model with regard to previous models that have been proposed. We also show that our model is consistent with several empirical observations of ant colonies.

## 1. Introduction

Darwinian natural selection explains how traits can evolve through time. However, the mechanisms involved behind the evolution of discrete novel traits are relatively less well understood, and often involve proximate mechanisms such as phenotypic or developmental plasticity [1]. Group selection theory, first proposed by Darwin, has increasingly been garnering interest [2], and recent advances have been made both through theoretical approaches [3] and the accumulation of empirical evidence [4, 5]. It has been proposed that since very few members of an ant colony reproduce, the colony as a whole may behave as a ‘superorganism’, with natural selection being shifted to the colony level [6, 7, 8]. Evidence for this is also seen through the existence of certain morphological forms that seem to have evolved through ecological specialization to benefit colony functioning (for example: specialized door-blocking castes in *Cephalotes* [9] and *Colobopsis* [10]).

Further, social Hymenopterans often have multiple morphologically and behaviorally distinct ‘castes’ within the same species. Individuals which have the same genetic code can show greatly differing phenotypes. The presence of such distinct castes is seen in ants, termites [6], bumblebees [11], aphids [12], thrips [13], and the clonal larvae of some parasitic wasps [14] and trematodes [15]. It is thus likely that the presence of castes has convergently evolved multiple times in social animals. However, the mechanism through which novel castes can evolve is unclear, since very few members of the colony reproduce and pass on their genes. Molet *et al*. have hypothesized that in ants, erratically observed ‘intercastes’, which are likely developmental recombinations of existing castes, may be the evolutionary precursors which evolve into novel castes [16, 17]. However, since intercastes may often initially be less functional than a member of any existing caste in the initial stages, one would naively never expect intercastes to persist in populations that are under natural selection at the individual level. Molet et.al. provided a verbal model to justify why such intercastes would not be wiped out by evolution, and how individuals that initially have low fitness could still persist in ant colonies[16].

The law of diminishing marginal utility is a well-known empirical law in economics and states that in any given system, the marginal utility of every additional good diminishes with the total number of goods. In this paper, we use this concept, along with related economic notions of productivity, consumption and marginal rate of substitution to model the evolution of monogynous eusocial ant colonies. In section (2), we lay out our assumptions and define quantities and concepts that are central to our model. This section drives most of the description in the subsequent sections. We then use this framework to provide a potential mechanism for the evolution of ant colonies through group selection in section 3. In the appendix, we show how various empirical observations of eusocial ant colonies are consistent with our model.

This paper thus formalizes a previously proposed verbal model, and develops a broad conceptual framework for a general group selectionist description of eusocial ant colonies.

## 2. The Model

### 2.1. The intuition

The meat of our argument relies on the fact that the costs and benefits of a single non-reproductive individual are likely negligible for survival of the colony. Ant colonies routinely lose workers to both biotic and abiotic factors, and this does not significantly hamper the functioning of the colony. However, if an individual were to somehow acquire a trait such as reproduction, that can be extremely beneficial to survival of the colony even when expressed in only a few members, then the colony benefits. Thus, colonies which produce a few ‘defective’ worker ants which do not contribute to colony functioning as much as a regular worker do not pay a severe price. On the other hand, if a colony can produce a few very useful ants, it is likely to greatly increase its own survival, and thus go on to produce more daughter colonies. Thus, if only a relatively small number of individuals in a colony undergo a mutation, the other members of the colony can buffer the entire colony from strong negative selection, acting somewhat analogous to an evolutionary capacitor.

### 2.2. Assumptions

#### Assumption 1.

We assume that every ant colony is founded by a single ‘queen’ ant, and attains maturity in finite time. Thus, our model only applies to monogynous colonies.

A colony is defined to be *mature* if it has a well-defined, time-independent size.

#### Assumption 2.

We assume that queens and males produced by these colonies disperse, mate only with ants from other colonies, and go on to produce their own, independent colony.

We define a *task* as any activity performed by an ant in a colony that benefits the colony in some way. For example: foraging, defense, and reproduction are all tasks.

#### Assumption 3.

We assume that the ability of an adult ant to perform a task is a function of its morphology, physiology, development, behavior, or a combination of all of these factors.

#### Assumption 4.

We assume that the types of offspring produced by a queen and the fractions of offspring that are of each type are independent heritable traits in queens and are subject to small mutations.

These assumptions directly imply that natural selection in our model acts at a ‘colony-level’. Colonies which produce more queens/males tend to produce more ‘daughter colonies’, and hence any trait carried by this colony will spread in the population. **Note:** The proximate mechanisms evolved in caste determination of Hymenopterans is still being investigated. Some species seem to show a genetic bias, and in others, caste seems to be determined entirely through epigenetic or developmental factors [18, 19, 20]. Regardless of the proximate cause, we assume only that the general pattern of caste determination is heritable. For example, if the determination is epigenetic or developmental, then the heritable information would be the molecular threshold of stimulation needed in order to determine the caste.

### 2.3. Utility Functions

Consider a mature colony of size *n* which needs to perform *η* tasks which are in the ordered set T = {*τ*_1_,…, *τ_η_*}. Different tasks have different purposes for the colony. Some tasks like foraging involve work done by ants by expending energy. Others, such as defense of the colony, protect the colony from potential energy losses (due to loss of colony members to predation). Thus, each ant either does some work, or ‘saves’ some work. We can quantify this (the work done/saved by the *i*th ant in doing the *j*th task) as a positive quantity *ω_ij_*. We define the *Usefulness* of the *i*th ant at performing the *j*th task as:

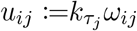

if the ant performs the *j*th task, and 0 if the ant does not perform the *j*th task. *ω* is a measure of how well an ant performs a given task (and can be empirically quantified in a variety of ways, with the particular metric probably being dependent on the task at hand), and *k_τ_j__* is a constant between 0 and 1 which quantifies the importance of the *j*th task for the functioning of the colony. This is the definition that we use for non-reproductive tasks. If the *j*th task is reproduction, we instead define the usefulness of the *i*th ant as:

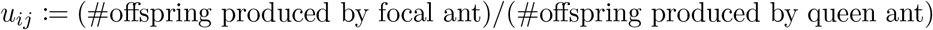

where the ‘queen ant’ is the ant that acted as the foundress of the mature colony.

We also define the *task specific energy consumption* of an individual ant as:

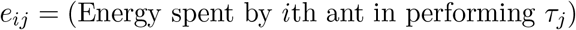

We define the *competence c_i_* of the *i*th ant as:

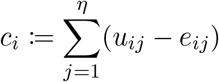

We also define the *utility* 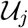 of a task *τ_j_* in the colony as:

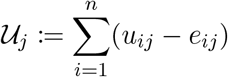

For large values of *n*, it may be useful to define the fraction of the population performing the task *i* as 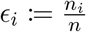. where *n_i_* is the total number of individuals in the colony with non-zero individual utility for the task *τ_i_* (*i.e*. the number of individuals which perform the task *τ_i_*). We define the *fractional marginal utility* as:

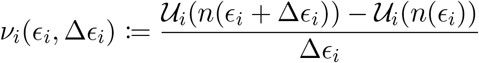

where Δ*ϵ_i_* is a small change in *ϵ_i_*. In the infinitesimal limit, we have:

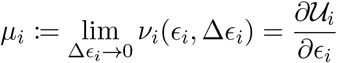

### 2.4. Productivity and consumption functions

We next define the *Gross Productivity function of a colony* as:

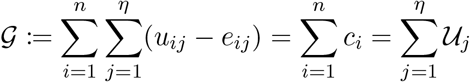

Thus, the competence of an individual is a measure of how much the presence of that individual contributes to the gross productivity of the colony, and the utility of a trait is a measure of how much the presence of the trait contributes to the gross productivity of the colony.

We define a *subsistence function* 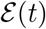 for the colony as:

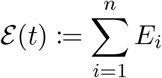

where *E_i_* is the energy that needs to be consumed by the *i*th ant of the colony in order to survive when it is not performing any tasks. The value of *E_i_* will depend on the morphology of the *i*th ant, and in practical terms, can be measured by determining the basal metabolic rate of the *i*th ant.

#### Assumption 5.

We assume that 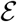 grows linearly with an increase in *n*, when *ϵ_i_* is held constant for all tasks.

This is a reasonable assumption to make, since each individual ant of a certain morphological type on average consumes the same amount of energy, and, if *ϵ_i_* is kept constant for all tasks, then, by consequence, the fraction of ants which have a particular morphology is also kept constant. We also define a Net Productivity function 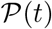 as follows:

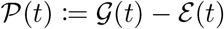

#### Assumption 6.

Crucially, we assume that given a colony of size *n*, the per-capita net productivity 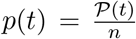 acts as a maximand for selection, *i.e*. natural selection tends to select colonies which have higher per-capita net productivity. Explicitly, if a colony has net productivity *p* and fitness *w*, we assume that:

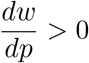

Alternately, given a colony of size *n*, net productivity 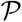,

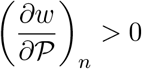

Note that here, by ‘fitness’, we refer to the absolute reproductive fitness of a colony, measured by the number of daughter colonies that it produces.

#### Assumption 7.

We assume that for any given task *τ_i_*, the utility function 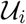 follows the law of diminishing marginal utility past a threshold both when varied with *n_i_* (keeping *ϵ_i_* constant for all tasks *τ_i_*), and when varying any given *ϵ_i_* (when *n* is kept constant), *i.e*.:

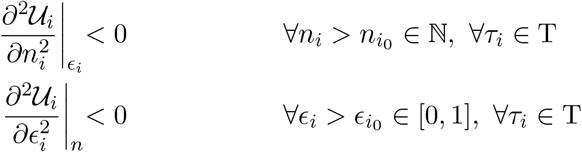

This implies that 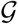 follows a similar behavior past a threshold i.e.

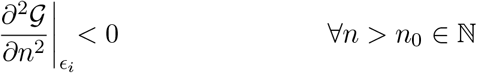

#### Assumption 8.

We further assume that the following trends hold:

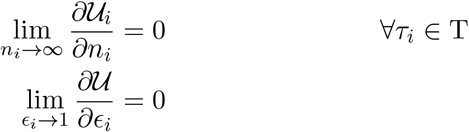

It is intuitively easy to see why assumptions (7, 8) are reasonable. They are justified by the fact that the resources around an ant colony are generally finite. Thus, if for example, we were looking at the individual usefulness of foragers, we would expect that indefinitely increasing the number of nest members or the number of foragers would not keep increasing the utility of foraging at a constant rate. In fact, as you keep adding foragers, you would expect the resources to become increasingly limited, leading to logistic-like growth of the utility functions. Since 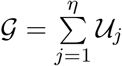, we have:

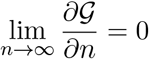

## 3. Results

### 3.1. Division of labour

Consider a mature colony of size *n*. Let the fraction of individuals which carry out a given task *τ_i_* be given by *ϵ_i_*. By assumption (6), we look for the condition at which each partial derivative 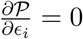.

We define the *Marginal Rate of Substitution* (MRS) *M_ij_* between two tasks as:

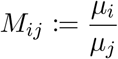

Where *μ_i_* and *μ_j_* are the fractional marginal utilities of the *i*th and *j*th task respectively. Rearranging, we get

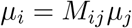

This is a comparison of the marginal utilities of task *τ_i_* and *τ_j_* and tells us that adding one more ant to perform task *τ_i_* is “*M_ij_* times as useful” as adding one ant to perform task *τ_j_* instead, in terms of contributions to the gross productivity of the colony.

By assumption (7), we see that as *ϵ_j_* keeps increasing (keeping *ϵ_i_* fixed), *μ_j_* keeps reducing. Thus, *M_ij_* increases, and it is more beneficial to the colony if ants start performing *τ_i_* instead. This is how division of labour evolves *i.e*. it is more useful if at any given time, different groups of ants perform different tasks, instead of all ants performing the same task.

It is evident that an equilibrium will be attained only when the marginal utility of both tasks is equal, *i.e* there can be no further increase in 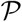 by individuals switching from one task to the other. This means that *M_ij_* = 1.

Extending this to *η* tasks, we see that the fixed proportions of workers of a colony that perform a given task can be determined by solving:

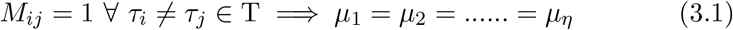

constrained to the condition:

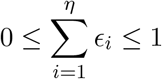

This condition is true because at any given instant of time, an ant can perform at most 1 task. Note that equation (3.1) is a constraint on the colony as a whole, and not on individuals. Therefore, an ant which engages in task *τ_i_* at some point of time can switch over to task *τ_j_* at some later point of time, as long as this switch is also accompanied by some other ants also switching tasks.

### 3.2. Redistribution of tasks helps the colony

Here we study what happens if a fraction of individuals who perform a particular task die. We show that in our system, a colony redistributes its workers such that this loss is supplemented by a readjustment of workers from other tasks.

Let us assume that some number of individuals which performed the task *τ_i_* died. Before any individuals died, the system was in equilibrium, and equation (3.1) held, **i.e*. M_ij_* = 1 ∀ *τ_j_* ≠ *τ_i_* ∈ T. Now, if *δϵ_i_* is the fraction of dead workers out of the initial *ϵ*_*i*0_, we see that:

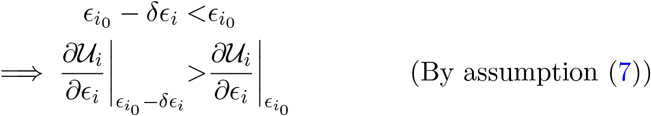

Thus, we see that *M_ij_* > 1 ∀*τ_j_* ≠ *τ_i_* ∈ T, and a ‘redistribution’ of workers such that more workers are allocated to perform *τ_i_* would increase the productivity of the colony. It is easily seen that even after this redistribution, the new net productivity is still less than the productivity of the colony before the deaths occurred. Thus, this type of redistribution only serves to ‘buffer’ the loss of the dead individuals.

### 3.3. Main proof

We define a caste as a morphologically and behaviorally specialised type of ant which performs a particular, fixed set of functions in the colony. We begin by examining our fitness function *w*(*t*) which measures the average number of daughter colonies produced by a given colony at a given point of time. If *w*(*t*) is greater than 1, a colony will not go extinct and the genes of its foundress are passed on in the population. We examine the rate of change of fitness,

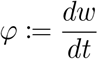

For the sake of simplicity, we will assume that we start with a mature colony that has only a queen and a single other caste, which we shall call the ‘worker’ caste. The same arguments proposed here hold regardless of the initial state of the colony, as will easily be seen.Consider a reference colony of size *n* which has a basal fitness of *w*_0_. Let a mutation occur in the daughter colony produced by this queen. By assumption (4) this mutation could affect either the types of non-reproductive offspring produced, the fraction of each of those types of offspring produced, or both. Let us assume both factors are affected, and ants which possess a previously unseen trait (or combination of traits) are now born in this colony. If this mutation increases the competence of these ants, it automatically increases the fitness of the colony, and this trait will spread in the population. Let us assume that this mutation decreases the fitness of the colony by negatively affecting the competence of the ants that are produced. Let the competence of the new ants be *c_n_*(*t*). It is clear that the rate of change of fitness will depend on the fraction of offspring who are affected. Let the number of such offspring produced at time *t* be *m*(*t*).

Now, at every instance in which the colony (or its descendants) breeds, there is some chance that there is another mutation which further modifies this trait. Thus, this trait is now effectively performing a random walk in mutation-space. Let *θ_−_* be the time elapsed before this trait becomes beneficial to the colony and the marginal net productivity becomes positive again. Qualitatively, this is the point at which ants which express the trait are ‘at least as useful’ as a worker. If, in this time, the fitness never dropped below 1, the trait is never removed and is successfully fixed in the population. Let us assume that the fitness drops below 1 before the trait becomes beneficial. From the point when fitness first started increasing again, let *θ_+_* be the time taken before the fitness grows back to 1.

Thus, the total time elapsed is *θ* = *θ_−_* + *θ_+_*, and the total change in fitness Δ*w* during this period will be given by:

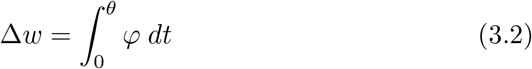

Now, this trait can be removed from the population at two time periods:

#### Case 1: Extinction during *θ_−_*

In the entire period of *θ_−_*, the fitness is decreasing throughout. Thus, if

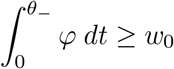

the trait will be removed from the population since the fitness of the colony is reduced to 0. Biologically, this either means that the trait was so detrimental that it drastically reduced the fitness of the colony in a short period of time, or that the trait either took too long to become beneficial to the colony, or was not expressed in high enough numbers to become beneficial to the colony.

#### Case 2: Extinction during *θ_+_*

Assuming that the population reaches the point where the trait becomes beneficial without going extinct, it can *still* go extinct if the fitness does not rise back up quickly enough. Thus, if the total number of colonies produced at the end of time *θ_−_* is *N_−_* and the time averaged fitness in subsequent time *ϕ* is given by 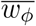, then the condition for extinction is that:

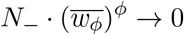

Let us assume that this, too, doesn’t happen, and the population successfully reaches a fitness of 1. We would like to know how to maximize the chances of reaching this stage.

We now draw on assumption (6) and assume that examining how productivity changes will allow us to draw inferences about how the fitness changes. We now have exactly two kinds of ants: Those that have the mutation, and those that don’t. We can safely assume that all the ants which have the trait are identical to each other, and that, likewise, all ants which do not have the trait are also identical to each other. Thus, our productivity can be split into two terms: Let *p_m_* be the per-capita productivity of the ants which possess the mutation, and let *p*_0_ be the per-capita productivity of the ants which do not possess the mutation (This is why we assumed a colony with only one kind of worker. If we do not, we will have several different *p*_0_ terms, but the basic arguments do not change). Now, we have:

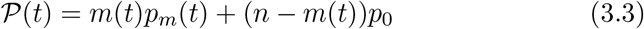

Differentiating equation (3.3) with respect to *m* gives us:

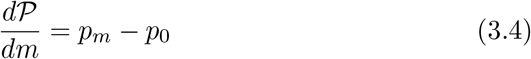

and differentiating equation (3.3) with respect to time gives us:

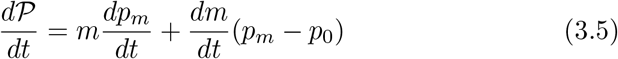

We now have the necessary equations, and can examine the optimal conditions at each phase of this process.

#### Phase 1

0 < *t* < *θ_−_*

During this period, the trait is not useful, and *p_m_* is less than *p_0_*. Thus, the quantity *p_m_* − *p*_0_ is *negative*. A cursory glance at equation (3.4) tells us that at this stage, 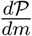 is a decreasing function. Thus, mutations that *minimize m* tend to maximize 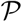 (and hence fitness), since we are assuming that *n* is constant. Examining equation (3.5) also tells us that either small positive values, or large negative values of 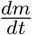 will tend to increase 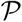. Thus, until the trait becomes beneficial, it should only affect a *small* proportion of the population.

#### Phase 2

*θ_−_* < *t* < *θ_+_*

During this period, the trait is useful. Thus, the quantity *p_m_* − *p*_0_ is *positive*. Examining equation (3.4), we see that 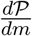 is now positive, and hence, increasing *m* will increase 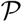 (and hence fitness). Additionally, examining equation (3.5) tells us that we must now try to maximize 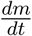. For as long as 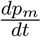 is positive (*i.e*. for as long as the trait becomes more and more useful with time), maximizing it is beneficial. Thus, in this phase, populations in which m and *p_m_* both increase rapidly will be less likely to go extinct, which is in line with what we expect intuitively.

In conclusion, we have shown that in our model, novel complex traits can evolve in ant colonies if the following conditions are satisfied:

i. The trait is not initially severely detrimental to the fitness of the colony.
ii. The trait can, when developed, in principle become useful to the colony even when expressed in small numbers.
iii. The trait is initially expressed in only a small number of the non-reproductive offspring, until it becomes useful.
iv. Once the trait becomes beneficial, its presence in the population increases rapidly.

In real life, ants which satisfy these conditions are the ‘intercastes’, which often have reduced utility, and are produced erratically in small numbers. In fact, both the soldier caste and the ergatoid queen caste exhibit the telltale signs of having developed from developmental mosaics of worker and queen castes [16]. Thus, our model justifies the conceptual model proposed by Molet *et al*.

Note that our model predicts that *m* should continually increase only until *p_m_* = *p*_0_ (i.e the *per capita* productivity of both types of ants is equal).

## 4. Discussion

Our model relies on several assumptions, some more realistic than others. Some of these assumptions may seem too idealistic. At this stage, we would like to point out that most mathematical models, due to the nature of model construction, must tread a fine line between idealism and realism. A model that is too realistic risks losing predictive power by having to incorporate too many parameters, and a model that relies on too many assumptions risks losing accuracy in the real world. Unfortunately, this is a limitation of using mathematics to make predictions about complex, real-world phenomena. We hope that we have managed to tread this fine line, in the sense that while some of our assumptions may not be very realistic, they may help provide a model that is a crude approximation of the real world. Our model is not intended to be right in all its gory details, it is only intended to be useful and/or insightful.

### 4.1. Comparisons with previous models

Molet *et al*. [16, 17] provided a verbal model for how novel castes may evolve in ant colonies even when such intercastes initially do not contribute to colony functioning. Our model mathematically shows that the verbal model that they propose is justified. Furthermore, it presents a very general conceptual framework for colony-level optimization theory. We are far from the first to present optimization models for eusocial colonies. Oster and Wilson developed an ergonomic theory to determine which castes should be maintained in a population of colonies [21, 22]. This was accomplished through a linear optimization model which lead to several predictions. The model that we present is a more general optimization model than that of Oster and Wilson, but some of the predictions that the two models make are the same. Both models predict that the ratio of castes that are present in a colony should be close to the optimum value. Though empirical evidence is relatively scarce, support for optimal caste ratio theory is seen in ants [23, 24, 6], termites [6], the clonal larvae of polyembryonic wasps [25], and the clonal larvae of trematodes [26, 27, 28, 29] (But see [30] for an example in which proportional task allocation follows seasonal schedules independent of colony demand, and are enforced by developmental constraints.)

Hasegawa [23] has developed an optimizatiom model on an *n*-dimensional hyperspace, and also provided empirical evidence for his model in the dimorphic ant species *Colobopsis nipponicus*. However, in Hasegawa’s model, the net ‘efficiency’ (which is equivalent to the net productivity function in our framework) is given by the product of the individual efficiencies. That is to say, if there are *τ* tasks in a colony, and if the state of the colony (given by the proportion of each caste in the colony) is *r*, then, in Hasegawa’s model, the total colony efficiency is given by 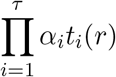, where *t_i_*(*r*) is the ‘efficiency function’ for the ith task, and *α_i_* is a quantity that determines the importance of the *i*th task for colony fitness. Thus, the quantity *α_i_t_i_*(*r*) is equivalent to the utility function 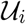 in our framework. Hasegawa uses the product of individual efficiency functions to arrive at the net efficiency. This means that a small number of ‘unimportant’ tasks (low *α_i_*, or, in our framework, low 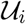) can heavily influence the net efficiency, which is unrealistic. We thus believe that our formulation is more robust to the existence of relatively unimportant tasks.

Rueffler *et al*. [31] develop a framework for optimization of the performance of functionally specialized ‘modules’ in organisms. This is analogous to our framework for ant colonies, since ant colonies can be viewed as superorganisms in which each caste is analogous to a module (see Appendix A.5). We also provide a rational justification from economic first principles for such an optimization.

The idea that eusocial insect societies are comparable to a superorganism, and thus may be subject to selection at a colony-level, is an old one [6, 7, 8]. The idea that small mutations may be buffered by the rest of the colony has also been proposed before [16]. The idea that caste ratios may be tuned to optimal values is not new either [21, 22]. We develop a single unifying framework that incorporates all of these phenomena, and includes these theories within it. Our model is also consistent with several other empirical observations (see Appendix).

### 4.2. Potential experiments to determine utility and productivity functions

Determining utility functions and productivity functions for ant colonies is no easy task, and will require elegant experiments, and quite possibly novel experimental techniques altogether. However, if these functions can be determined empirically, our model should provide a framework to predict the future direction of evolution that is likely to be followed by this colony in a given fixed environment. Our model also makes one very general prediction that should be true of all colonies: We propose that the model can be tested by experimentally generating ‘knock out’ colonies (the term being borrowed from genetics), by selectively eliminating all members of a given caste from a colony of ants, and then eliminating all subsequent larvae that mature into the focal caste. Set up with the appropriate controls, this will allow us to look for ‘task redistribution’, whereby some ants move away from their task to perform the task that the eliminated caste used to perform. This is a natural prediction of our model (independent of the form of productivity and utility functions). The presence of task redistribution would lend credibility to our model, and the absence of any redistribution, even among morphologically similar castes, would falsify it. Furthermore, if we had access to a large enough number of colonies so as to generate a sizable number of knockout colonies for each caste present in the species, we would be able to directly measure utility functions of these castes by monitoring colony functioning for each class of knockout colony. Specific measures of utility and productivity functions will allow for very specific predictions about quantities such as colony size, caste ratios in a colony, and number of castes.

### 4.3. Proxies for Utility and Productivity functions

For a monomorphic colony of size *n* performing *τ* tasks, we can write the gross productivity function as 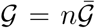, and the total energy consumed as 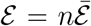. For such a colony, assuming workers are identical, from equation 3.1, we know that:

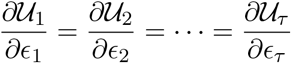

Let the per-capita utility of the *i*th task be given by 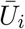. Then,

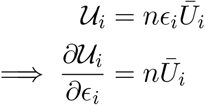

Since we know that the fractional marginal utilities of every task must be equal, we have 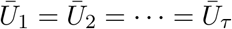, which means that the per-capita utility is the same for all tasks. Since all ants are assumed to be morphologically identical, the total mass of all ants performing any given task can thus be used a proxy for the utility 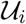 of that task, *i.e*.

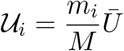

where *m_i_* is the total mass of all ants that are engaged in task *τ_i_*, and *M* is the total biomass of the colony. Since the sum of the utility functions over all tasks equals the gross productivity function, the biomass of the colony can be used as a proxy for the gross productivity function.

## Acknowledgements

We would like to thank Dr. Sutirth Dey and Dr. Raghavendra Gadagkar for their valuable comments on the manuscript. We would also like to thank two anonymous reviewers, whose comments greatly improved our manuscript.

## Appendix A. predictions and empirical observations that are consistent with our model

### Appendix A.1. The total number of castes has an upper bound

Our model shows that as the number of existing castes in a colony increases, the probability of yet another novel caste evolving decreases. Let there be *χ* castes in a colony, and let 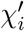 be the number of castes which have the ability to perform a task *τ_i_*, and let 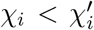 be the number of castes which actually perform this task. Mutations affect individuals as described in 3. We assume the worst possible case, and say that until time *θ_−_* (as defined in 3) has passed, the utility of mutated ants is equal to that of a dead ant. If some fraction 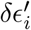 of a particular caste which actually performs *τ_i_* is affected by this mutation, we would expect redistribution of workers from the total number of ants from one or more of the 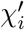 castes which are able to perform *τ_i_*. From this, we can draw two inferences: Firstly, since castes are generally specialized in terms of morphology and behavior (and thus the functions they perform), an increased number of castes will likely cause a reduction in the number of castes that can perform a task. Thus it is reasonable to make the following assumption:

#### Assumption 9.

As the number of castes in a colony increases, the number of individuals which can perform a given task is non-increasing, *i.e*.

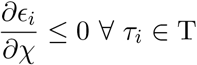

Under this assumption, it is clear that the effectiveness of the buffering as described in 3.2 is non-increasing as the number of castes increases, and thus, if a colony has more pre-existing specialized castes, the detrimental effect on the fitness of the colony during the *θ_−_* period is stronger, and the population is more likely to go extinct before the trait can become useful. In the event where certain tasks are essential for sustenance of the colony, a minimum utility may be essential for the survival of the colony, in which case, the population will survive only if the utility of that task exceeds a minimum threshold 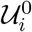. Thus, the condition to be met after redistribution is:

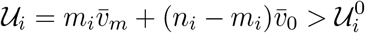

where *m_i_* is the number of mutants, *n_i_* = *ϵ_i_n* is the total number of ants which can perform *τ_i_*, and 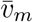 and 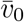 are the per-capita utilities of the mutants and non-mutants respectively. Differentiating with respect to *ϵ_i_*, we get:

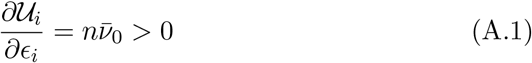

But as a consequence of assumption (9), we see that:

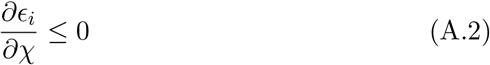

and thus,

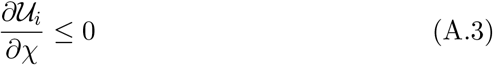

Equation (A.3) tells us that as the number of castes in a colony increases, 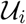 is non-increasing, and thus, it becomes harder to keep the utility of these tasks above the minimum utility required for colony maintenance.

### Appendix A.2. Variation of number of castes with colony size

In section 3, we illustrated the mechanism through which novel castes can evolve. Integral to this mechanism is the ‘buffering capacity’ of nonmutated ants which can prevent the colony from going extinct before the trait becomes beneficial. It is clear that the buffering capacity is higher if the colony includes more non-mutated ants in the initial stages of evolution. Thus all other things being equal, we would expect species with larger colony sizes to have a stronger buffering capacity and thus it is more probable for a species with a larger colony size to evolve a new caste. This is in line with empirical observations [32, 33].

### Appendix A.3. Variation in gyne to worker ratio with colony size

Schmidt *et al*. reported that in the Pharoah ant (*Monomorium pharaonis*), a species that is polygynous and exhibits extensive intranidal mating, the gyne to worker ratio decreases as colony size increases [34]. In a completely intranidal colony, since a single male can fertilize multiple gynes, it would be fair to approximate the scaling of the number of offspring produced by a colony to be linear with respect to the number of gynes. Thus, for a colony of size *n* and *n_g_* gynes, if we assume generations are discrete in time, then, at any point *t*, we can write:

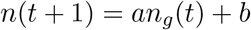

where *a* and *b* are positive constants, and the generation time is taken to be 1 for simplicity. Thus, an increase in the number of gynes will lead to a linear increase in colony size *n*. Now, for optimal functioning of a colony, we assume that the net productivity 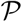 must be maximized, or, given the colony size *n*, that the per-capita productivity *p* must be maximized. We know that:

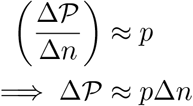

Thus, a bigger colony will require a proportionally bigger net productivity 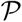 to be able to sustain itself at optimal levels. However, because of the law of diminishing marginal utility, we know that we will need a non-linear increase in the number of workers to achieve the same linear increase in net productivity, and thus the number of workers will have to be several times more in the bigger colony to be able to sustain it. Thus, we expect the gyne to worker ratio to decrease as colony size increases, which lines up with the observations of Schmidt *et al*.[34].

### Appendix A.4. Variation of number of castes with latitude

Wilson [6, 22] pointed out that species which live in tropical climates tend to show much higher levels of caste diversity and specialization than species found in temperate climates. Our model allows for this as well. In section 3, we showed how our model allows for the evolution of novel castes. We also highlighted that if the mutated trait proves to be useful, selection pressure would lead to those colonies being selected which have that trait. If we consider a utility function 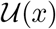 as a function of the value of a trait *x*, we would expect selection pressure to eventually move the system towards a state with greater 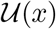, and thus, natural selection would select the colony which had ants that displayed the phenotype at which 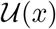 has a local maximum. This introduces a directionality in the evolution of morphological variations. However, as we saw earlier, for any given novel (individual-level) trait, there is an initial period when the trait is not useful for the colony, and mutated ants thus pose an additional cost to the colony. However, since the magnitude of benefit is dependent on the utility function, variation in the utility function will lead to variation in the utility of the phenotype. Thus, the directionality that had been induced is lost if the utility function varies, and it is unlikely that the trait reaches the point where it becomes beneficial before being wiped out in the manner described in section 3. Note that the utility function of any trait is strongly dependent on the environmental conditions, and thus, in temperate climates, which experience greater fluctuations in abiotic factors as seasons change, we would expect fewer castes to be present, and that those castes which are present are generalists (*i.e*. are not too phenotypically different from the workers) and not specialists, which is in line with Wilson’s observation.

### Appendix A.5. Multicellularity

The framework that we propose is a general one, based on basic economic principles, and can be applied to biological systems at various levels. While this document focuses on ant colonies, for the purposes of our model, eusocial insect colonies and primitive multicellular organisms are functionally equivalent. A multicellular organism behaves very similar to a colony: All the cells carry the same genes; only a few cells reproduce, and the rest serve to maintain optimal functioning of the organism. Thus, we make the bold claim that the same arguments proposed here also serve to explain how differentiation could evolve in multicellular organisms, with the ‘castes’ in that case referring to the number of cell types, as in [31]. It has been widely observed that the number of cell types tends to increase as the number of cells increases [32, 35, 31]. In the context of our model, this makes sense, since an increase in the number of cells corresponds to a larger buffering effect, and hence a higher likelihood of evolution of novel cell types.

## References

[1] M. J. West-Eberhard, Developmental plasticity and the origin of species differences, Proceedings of the National Academy of Sciences 102 (suppl 1) (2005) 6543–6549.

[2] M. E. Borrello, The rise, fall and resurrection of group selection, Endeavour 29 (1) (2005) 43–47.

[3] A. Gardner, A. Grafen, Capturing the superorganism: a formal theory of group adaptation, Journal of evolutionary biology 22 (4) (2009) 659–671.

[4] C. Hou, M. Kaspari, H. B. Vander Zanden, J. F. Gillooly, Energetic basis of colonial living in social insects, Proceedings of the National Academy of Sciences 107 (8) (2010) 3634–3638.

[5] J. F. Gillooly, C. Hou, M. Kaspari, Eusocial insects as superorganisms: insights from metabolic theory, Communicative & integrative biology 3 (4) (2010) 360–362.

[6] E. O. Wilson, Sociobiology, the new synthesis, Harvard University Press, 1975.

[7] D. S. Wilson, E. O. Wilson, Rethinking the theoretical foundation of sociobiology, The Quarterly review of biology 82 (4) (2007) 327–348.

[8] H. K. Reeve, B. Hölldobler, The emergence of a superorganism through intergroup competition, Proceedings of the National Academy of Sciences 104 (23) (2007) 9736–9740.

[9] S. Powell, Ecological specialization and the evolution of a specialized caste in cephalotes ants, Functional Ecology 22 (5) (2008) 902–911.

[10] H. Fujioka, M. S. Abe, Y. Okada, Observation of plugging behaviour reveals entrance-guarding schedule of morphologically specialized caste in colobopsis nipponicus, Ethology 125 (8) (2019) 526–534.

[11] J. Peat, J. Tucker, D. Goulson, Does intraspecific size variation in bumblebees allow colonies to efficiently exploit different flowers?, Ecological Entomology 30 (2) (2005) 176–181.

[12] D. L. Stern, W. A. Foster, The evolution of soldiers in aphids, Biological Reviews 71 (1) (1996) 27–79.

[13] B. J. Crespi, Eusociality in australian gall thrips, Nature 359 (6397) (1992) 724–726.

[14] Y. Cruz, A sterile defender morph in a polyembryonic hymenopterous parasite, Nature 294 (5840) (1981) 446–447.

[15] R. F. Hechinger, A. C. Wood, A. M. Kuris, Social organization in a flatworm: trematode parasites form soldier and reproductive castes, Proceedings of the Royal Society B: Biological Sciences 278 (1706) (2011) 656–665.

[16] M. Molet, D. E. Wheeler, C. Peeters, Evolution of novel mosaic castes in ants: modularity, phenotypic plasticity, and colonial buffering, The American Naturalist 180 (3) (2012) 328–341.

[17] S. Londe, T. Monnin, R. Cornette, V. Debat, B. L. Fisher, M. Molet, Phenotypic plasticity and modularity allow for the production of novel mosaic phenotypes in ants, Evodevo 6 (1) (2015) 1–15.

[18] E. Abouheif, Ant caste evo-devo: it’s not all about size, Trends in Ecology & Evolution 36 (8) (2021) 668–670.

[19] A. M. Rafiqi, A. Rajakumar, E. Abouheif, Origin and elaboration of a major evolutionary transition in individuality, Nature 585 (7824) (2020) 239–244.

[20] W. Trible, D. J. Kronauer, Ant caste evo-devo: size predicts caste (almost) perfectly, Trends in Ecology & Evolution 36 (8) (2021) 671–673.

[21] G. F. Oster, E. O. Wilson, Caste and ecology in the social insects, Princeton University Press, 1978.

[22] E. O. Wilson, The ergonomics of caste in the social insects, The American Naturalist 102 (923) (1968) 41–66.

[23] E. Hasegawa, The optimal caste ratio in polymorphic ants: estimation and empirical evidence, The American Naturalist 149 (4) (1997) 706–722.

[24] L. Passera, E. Roncin, B. Kaufmann, L. Keller, Increased soldier production in ant colonies exposed to intraspecific competition, Nature 379 (6566) (1996) 630–631.

[25] J. A. Harvey, L. S. Corley, M. R. Strand, Competition induces adaptive shifts in caste ratios of a polyembryonic wasp, Nature 406 (6792) (2000) 183–186.

[26] T. Kamiya, R. Poulin, Caste ratios affect the reproductive output of social trematode colonies, Journal of evolutionary biology 26 (3) (2013) 509–516.

[27] M. M. Lloyd, R. Poulin, Reproduction and caste ratios under stress in trematode colonies with a division of labour, Parasitology 140 (7) (2013) 825–832.

[28] M. M. Lloyd, R. Poulin, Geographic variation in caste ratio of trematode colonies with a division of labour reflect local adaptation, Parasitology research 113 (7) (2014) 2593–2602.

[29] C. MacLeod, R. Poulin, C. Lagrue, Save your host, save yourself? caste-ratio adjustment in a parasite with division of labor and snail host survival following shell damage, Ecology and evolution 8 (3) (2018) 1615–1625.

[30] C. L. Kwapich, W. R. Tschinkel, Demography, demand, death, and the seasonal allocation of labor in the florida harvester ant (pogonomyrmex badius), Behavioral Ecology and Sociobiology 67 (12) (2013) 2011–2027.

[31] C. Rueffler, J. Hermisson, G. P. Wagner, Evolution of functional specialization and division of labor, Proceedings of the National Academy of Sciences 109 (6) (2012) E326–E335.

[32] Y. Iwasa, S. Yamaguchi, Task allocation in a cooperative society: specialized castes or age-dependent switching among ant workers, Scientific reports 10 (1) (2020) 1–9.

[33] H. Ferguson-Gow, S. Sumner, A. F. Bourke, K. E. Jones, Colony size predicts division of labour in attine ants, Proceedings of the Royal Society B: Biological Sciences 281 (1793) (2014) 20141411.

[34] A. M. Schmidt, T. A. Linksvayer, J. J. Boomsma, J. S. Pedersen, Queen–worker caste ratio depends on colony size in the pharaoh ant (monomorium pharaonis), Insectes sociaux 58 (2) (2011) 139–144.

[35] J. T. Bonner, The evolution of complexity by means of natural selection, Princeton University Press, 1988.

